# A new counter-intuitive therapy for adult amblyopia

**DOI:** 10.1101/360420

**Authors:** Lunghi Claudia, Sframeli Angela Tindara, Lepri Antonio, Lepri Martina, Lisi Domenico, Sale Alessandro, Morrone Maria Concetta

**Author notes:** Co-senior authors (Sale Alessandro, Morrone Maria Concetta). Corresponding Author: Alessandro Sale, Neuroscience Institute, National Research Council (CNR), Via Moruzzi 1, 56124, Pisa, Italy.

## Abstract

Visual cortex plasticity is high during a critical period of early postnatal development, but rapidly diminishes with the transition to adulthood. Accordingly, visual disorders such as amblyopia (lazy eye), can be treated early in life by long-term occlusion of the non-amblyopic eye, but may become irreversible in adults, because of the decline in brain plasticity. Here we show that a novel counter-intuitive approach can promote the recovery of visual function in adult amblyopic patients: short-term occlusion of the amblyopic (not the fellow) eye, combined with physical exercise (cycling). After six brief (2h) training sessions, visual acuity improved in all ten patients (0.15±0.02 LogMar), and six of them also recovered stereopsis. The improvement was preserved for up to one year after training. A control experiment revealed that physical activity was crucial for the recovery of visual acuity and stereopsis. Thus, we propose a non-invasive therapeutic strategy for adult human amblyopia based an inverse-occlusion and physical exercise procedure.

## Introduction

Amblyopia is a neurodevelopmental disorder of vision^1,2^ with a prevalence ranging from 1 to 5% depending on the population tested^3^. The most frequent causes of amblyopia are strabismus, anisometropia and deprivation (for example due to cataracts) early in life^1,2^, within the so-called *critical period*, defined as the temporal window during development where the plastic potential of the visual cortex is maximal^4,5^. If the retinal image is degraded during development, the visual cortex disregards the eye providing degraded input and responds primarily to the eye providing the better signal. Amblyopia can be ameliorated in young children by the long-term occlusion of the better (non-amblyopic) eye^3^. However, in humans, treatment after the closure of the critical period dramatically decreases therapy efficacy requiring extensive occlusion of the non-amblyopic eye to produce only a modest improve of visual acuity (234 hours of occlusion per 0.1 LogMar acuity improvement^6^). New therapeutic strategies recently proposed for adult amblyopia involve either very prolonged visual training^7–12^, or invasive manipulations^13,14^, rendering them almost ineffective for clinical purposes.

Recently, new hope has emerged from evidence that the early visual cortex retains some degree of plasticity in adult humans^15–23^. Many studies have demonstrated a plastic response to visual deprivation: a few hours^15,16^ or a few days^17,18^ of binocular deprivation modulate excitability of the primary visual cortex. But also monocular occlusion of one eye for a few hours paradoxically boosts the deprived eye signal, shifting ocular dominance in favor of the deprived eye^19–22^. This counter-intuitive effect of monocular deprivation reflects a compensatory reaction of the visual cortex to deprivation aimed at maintaining the average cortical activity constant, indicating that some form of homeostatic plasticity^24^ is also present in adult humans. Importantly, this plasticity is mediated by a decrease of GABA concentration in the primary visual cortex^25^ and is enhanced by physical exercise^26^.

The modulatory effect of physical activity on visual plasticity is corroborated by several recent studies on animal models^27–30^: in adult rats and mice physical activity boosts visual cortical plasticity by modulating the levels of GABAergic inhibition^27–29^ via a specific somatosensory- visual circuitry^27–30^. Interestingly, visual cortical GABAergic interneurons are implicated in the mechanisms controlling the onset and offset of the visual critical period in these animal models^31^.

While physical exercise appears as a potentially suitable and non-invasive therapeutic strategy for adult amblyopia, no attempts have been made, to date, to investigate its effects on amblyopic human subjects. Focusing on adult anisometropic patients, we performed a counterintuitive experiment which combined short-term deprivation of the amblyopic eye with physical exercise. We found that after six short training sessions (2h each), visual acuity and stereopsis improved in adult patients, and that the improvement persisted for up to one year after the end of the treatment.

## Methods

### Subjects

Adult anisometropic patients were enrolled for the study after an ophthalmic screening examination in which detailed ocular and systemic anamnesis has been investigated (see Procedures section). Patients with a history of chronic ocular diseases other than amblyopia have been excluded from the study as long as patients presenting any chronic systemic disease. Similarly, patients with neurological or psychiatric disorder or under medication of any sort were excluded.

Out of 40 patients screened, 10 met the inclusion criteria and were enrolled for the study (four males, mean age 33±1.5, Clinical data reported in Table 1).

**Table 1.** Clinical Data.

| Patient ID | Age/sex | Refractive Error |  | Visual Acuity (LogMar) |  | Stereoaucity (ArcSec) | Fixation AE | Treatment history |
| --- | --- | --- | --- | --- | --- | --- | --- | --- |
|  |  | Right Eye | Left Eye | Right Eye | Left Eye |  |  |  |
| S1 | 28/F | + 0.25 +0.25x75 | +2.25 +2.25x55 | -0.06 | 0.62 | F | Central unsteady | Patched at age 5 |
| S2 | 29/M | +0.00 | -1.00x60 | -0.2 | 0.16 | F | Central | No Treatment |
| S3 | 27/F | -6.50 -3.00x10 | -0.25 -0.75x165 | 0.1 | -0.14 | 600 | Central | Patched at age 11 |
| S4 | 36/F | +0.25 -7.00x10 | -1.00 | 0.2 | -0.2 | F | Central | No Treatment |
| S5 | 35/M | +0.00 | +3.00x90 | -0.2 | 0.54 | 240 | Central unsteady | No Treatment |
| S7 | 34/F | -0.25x180 | +2.00 -1.50x180 | -0.1 | 0.78 | F | Central | Patched at ages 7-10 |
| S8 | 38/F | -3.25 -1.50x10 | -1.00 -2.75x180 | 0.0 | 0.46 | 600 | Central | Patched at age 5 |
| S9 | 40/M | +0.00 | +2.00 | -0.2 | 0.14 | 240 | Central | Patched at age 5 |
| S10 | 34/M | +0.50 | -0.50 -0.50x90 | 0.38 | -0.2 | 240 | Central | Patched at ages 7-13 |
| S11 | 28/F | +5.00 +1.00x80 | +0.00 | 0.52 | -0.1 | 600 | Central unsteady | No Treatment |

### Ethical Statement

Experimental procedures are in line with the declaration of Helsinki and were approved by the regional ethics committee [Comitato Etico Pediatrico Regionale—Azienda Ospedaliero-Universitaria Meyer—Firenze (FI)], under the protocol “Plasticità del sistema visivo” (3/2011).

### Apparatus and Stimuli

#### Binocular Rivalry

The experiment took place in a dark and quiet room. Visual stimuli were generated by the ViSaGe stimulus generator (CRS, Cambridge Research Systems), housed in a PC (Dell) controlled by Matlab programs. Visual stimuli were two Gaussian-vignetted sinusoidal gratings (Gabor Patches), oriented either 45° clockwise or counterclockwise (size: 2σ = 2°, spatial frequency: 2 cpd), presented on a uniform background (luminance: 37.4 cd/m2, C.I.E: 0.442 0.537) in central vision with a central black fixation point and a common squared frame to facilitate dichoptic fusion. Visual stimuli were displayed on a 20-inch Clinton Monoray (Richardson Electronics Ltd., LaFox, IL) monochrome monitor, driven at a resolution of 1024×600 pixels, with a refresh rate of 120 Hz. Observers viewed the display at a distance of 57 cm through CRS Ferro-Magnetic shutter goggles that occluded alternately one of the two eyes each frame. Responses were recorded through the computer keyboard.

### Procedures

#### Screening Examination

Uncorrected visual acuity (UCVA, ETDRS charts), best corrected visual acuity (BCVA, ETDRS charts, average of three different charts), intra-ocular pressure (IOP), color vision (evaluated with the Ishihara tables) and stereopsis (evaluated with TNO and Lang tests 1 and 2) have been evaluated and registered. All patients have also been subjected to pharmacological cycloplegia (1 eye drop of 1% Tropicamide administered three times with an interval of 5 minutes between the administrations) in order to register cycloplegic ametropia with an autorefractometer (Topcon RM8900, Topcon, Japan) and skiascopy (Heine Beta 200, Heine, Germany) after 20 minutes from last eye drop administration. During the screening visit, all patients underwent non-contact ocular biometry (IOLMaster 500, Carl Zeiss, Germany) and axial length (AL), anterior chamber depth (ACD) and white to white signal (WTW) have been registered. A complete examination of ocular motility has also been performed in order to exclude muscular causes of amblyopia: cover and uncover far and near tests, versions and objective convergence evaluation, Irvine test and 8 diopters Paliaga test for microstrabimus. Fixational quality was also assessed using microperimetry (Nidek MP-1 Professional). The screening examination was completed with endothelial cell count (Tomey EM 3000, Tomey, Germany), corneal topography (Sirius system, CSO, Firenze, Italy) to study keratometric parameters, slit lamp biomicroscopy (SL 9900, CSO, Italy) and complete phundus examination with both 90D and 20D lenses have also been performed.

After the aforementioned examinations, subjects presenting any ophthalmic pathology other than anisometropic amblyopia have been excluded from the study. Subjects presenting abnormal or unclear ocular findings (example.g., elevated IOP, presence of epiretinal membranes, low endothelial cell count, etc.) have been excluded from the study, as long as subjects presenting abnormal ocular motility that may have caused amblyopia.

BCVA (ETDRS charts) and Stereoacuity (TNO test) were also measured in the Follow Up tests performed one month, three months and one year after the end of the training procedure.

### Training procedure

#### Main experiment

Each training session lasted 2h and consisted in the combination of amblyopic eye occlusion and physical exercise. Occlusion of the amblyopic eye was performed using eyepatching. The eye-patch was custom-made of a translucent plastic material that allowed light to reach the retina (attenuation 15%) but completely prevented pattern vision, as assessed by the Fourier transform of a natural world image seen through the eye-patch. During the 2h of monocular occlusion, patients sat on a stationary bike equipped with a chair and a computer monitoring physical activity parameters (cycling speed, distance) and heart rate through a wireless chest band. Patients were instructed to cycle intermittently maintaining a heart rate between 110 and 120 bpm for 10 minutes, interleaved with 10 minutes rest; the experimenter controlled that physical exercise was performed according to these parameters. A 20’’ monitor (LG) was placed in front of the bike at a distance of 90 cm, patients watched a movie projected onto the monitor during the training. Before and after each training session, visual acuity (ETDRS charts), stereoacuity (TNO test) and sensory eye dominance (binocular rivalry) were assessed for each patient. This training was repeated six times over a four-week period: on three consecutive days during the first week of training and one day per week during the following three weeks (Figure 1).

**Figure 1.**
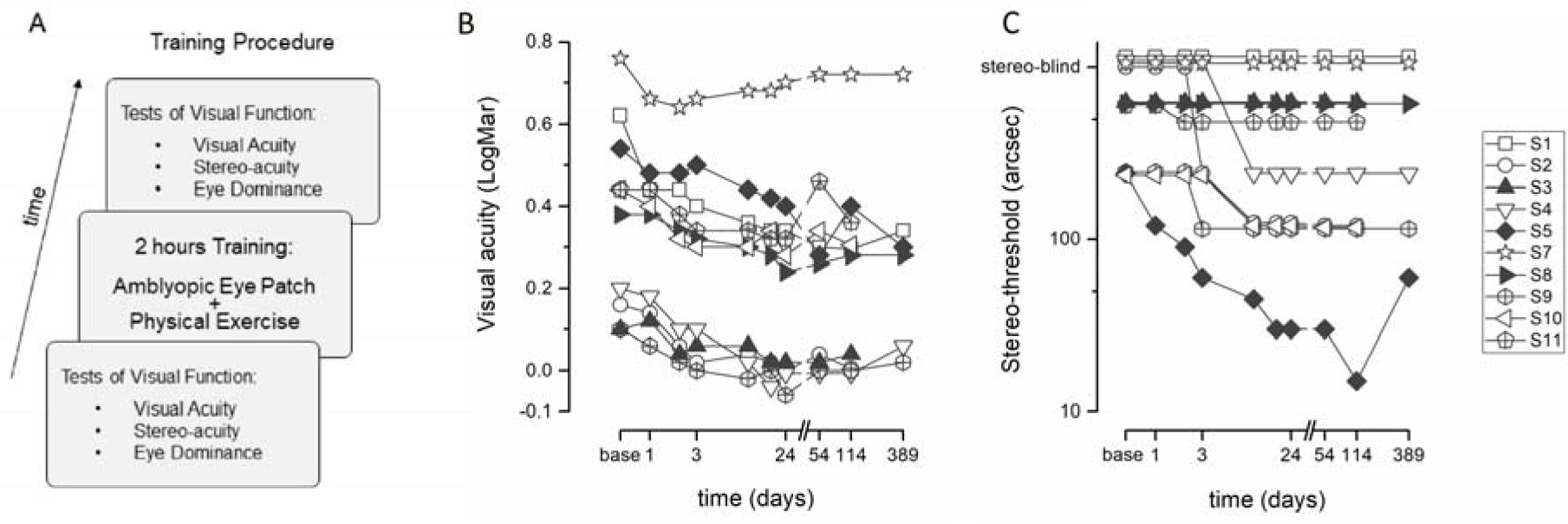
Training induces a recovery of visual acuity and stereo-threshold. (A) 10 adult anisometropic amblyopes underwent a training paradigm in which short-term (2h) occlusion of the amblyopic eye was combined with physical exercise (intermittent cycling on a stationary bike, 10 minutes of activity and 10 minutes rest interleaved, heartrate between 110 and 120 bpm). During the 2h training a movie was displayed in front of the stationary bike. Before and after each training session, sensory eye dominance (binocular rivalry between orthogonal gratings), visual acuity (ETDRS charts) and Stereo-thresholds (TNO test) were assessed for each patient. Visual acuity and stereo-thresholds were also assessed in follow-up measurements one, three and twelve months after the end of training. Each patient performed six 2h training sessions: three on consecutive days during the first week of training, and one session per week during the following three weeks. (B) LogMar visual acuity plotted as a function of time from the beginning of training. To achieve a robust quantification of visual acuity, each point represents the average of three different ETDRS charts. Different symbols represent the different performances after each training session. Data points after the break in the abscissa represent visual acuity obtained in follow-up measurements performed one 54 days, 124 days and 395 days after the first training session. (C) Stereo-thresholds plotted as a function of time from the beginning of training. Stereo-thresholds were obtained using the TNO test. For subjects showing course stereopsis with the TNO test (S3, S8, S11), stereo-thresholds were obtained using the LANG stereo-test.

#### Control Experiment

Each training session lasted 2h and consisted in the occlusion of the amblyopic eye (same procedure as main experiment) without simultaneous physical exercise. During the 2h of monocular occlusion patients sat at a distance of 90 cm in front of a 20’’ monitor (LG) and watched a movie. This training was repeated three times in three consecutive days. Before and after each training session, visual acuity (ETDRS charts), stereoacuity (TNO test) and sensory eye dominance (binocular rivalry) were assessed for each patient.

#### Binocular Rivalry

Each binocular rivalry experimental block lasted 180 seconds. After an acoustic signal (beep), the binocular rivalry stimuli appeared. Subjects reported their perception (clockwise, counterclockwise or mixed) by continuously pressing with the right hand one of three keys (left, right and down arrows) of the computer keyboard. At each experimental block, the orientation associated to each eye was randomly varied so that neither subject nor experimenter knew which stimulus was associated with which eye until the end of the session, when it was verified visually. Because of anisometropia, when rivalrous stimuli having equal contrast were presented, amblyopic patients do not report perceptual alternation, as vision is completely dominated by the non-amblyopic eye. In order to adjust the monocular stimuli contrast to compensate for the patients’ anisometropia, before the first training session, a few 60-seconds preliminary experimental blocks were performed. At each block, contrast of monocular stimuli was varied, increasing contrast of stimuli presented to the amblyopic eye (maximum contrast: 100%) and decreasing contrast of the stimulus presented to the non-amblyopic eye (minimum contrast: 20%), aimed at inducing perceptual alternations and equal predominance (if possible) of the two eyes. By performing this contrast adjustment procedure, binocular rivalry could be measured in six of the ten patients tested. In these six subjects, two binocular rivalry experimental blocks were acquired before and after each training session. The monocular contrast assessed in the preliminary test was kept constant throughout the experiment.

### Analyses

#### Binocular Rivalry

The perceptual reports recorded through the computer keyboard were analyzed using Matlab. Periods of mixed perception were excluded from the analyses. Mean phase durations (the average perceptual duration of each rivalrous stimulus) were computed for the two eyes. To quantify sensory eye dominance we obtained an index of ocular dominance, ranging from −1 (complete dominance of the non-amblyopic eye) to 1 (complete dominance of the amblyopic eye), by computing the contrast between the mean phase duration of the two eyes:

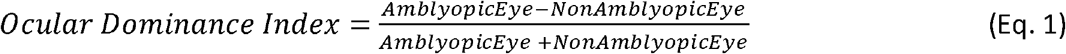

#### Statistics

Statistical analyses were performed using SPSS20 and Matlab softwares. BCVA, Stereoacuity and Sensory Eye Dominance before, during and after training were compared using Repeated Measures ANOVA (Greenhouse-Geisser correction was applied when the assumption of sphericity was violated) and paired-sample t-tests (α error fixed at 0.05). Correlations were computed using the Spearman’s correlation coefficient (rho), statistical significance assessed using a permutation test. The effect of training on visual acuity in the main and control experiment was compared using a within-subjects bootstrap sign test (10000 repetitions).

## Results

Ten adult anisometropic amblyopes (Table 1), selected for good fixation and with no other neurological deficit, underwent six brief training sessions over a four-week period (three on consecutive days during the first week and one session per week in the following three weeks: Figure 1). To assure a high homogeneity of the sample, only purely anisometropic amblyopes were included, discarding all patients showing microstrabismus. In each session, the amblyopic eye was occluded for two hours, during which patients intermittently cycled on a stationary bike while watching a movie. During the training period, visual acuity (ETDRS charts) significantly improved in all patients (Figure 1B), with a first improvement evident after the first two hours of patching (t(9)=2.37, p=0.04), increasing after each session. The average improvement after the sixth (and last) training session was 0.15±0.02 LogMar (t(9)=7.7, p<0.001, Figure 2A, Repeated Measures ANOVA, F(6, 54)=24.9, p<0.001, η^2^=0.73). This improvement was maintained in the follow-up measurements (Figure 3) obtained one month (t(9)=4.04, p=0.003), three months (t(9)=5.22, p<0.001) and one year after the end of training (t(5)=3.81, p=0.012). The initial visual acuity of the amblyopic eye and the acuity improvement observed at the end of the training were not correlated (Spearman’s rho=-0.16, p=0.67, Figure 4A), indicating that the effect of the training was independent of anisometropia severity.

**Figure 2.**
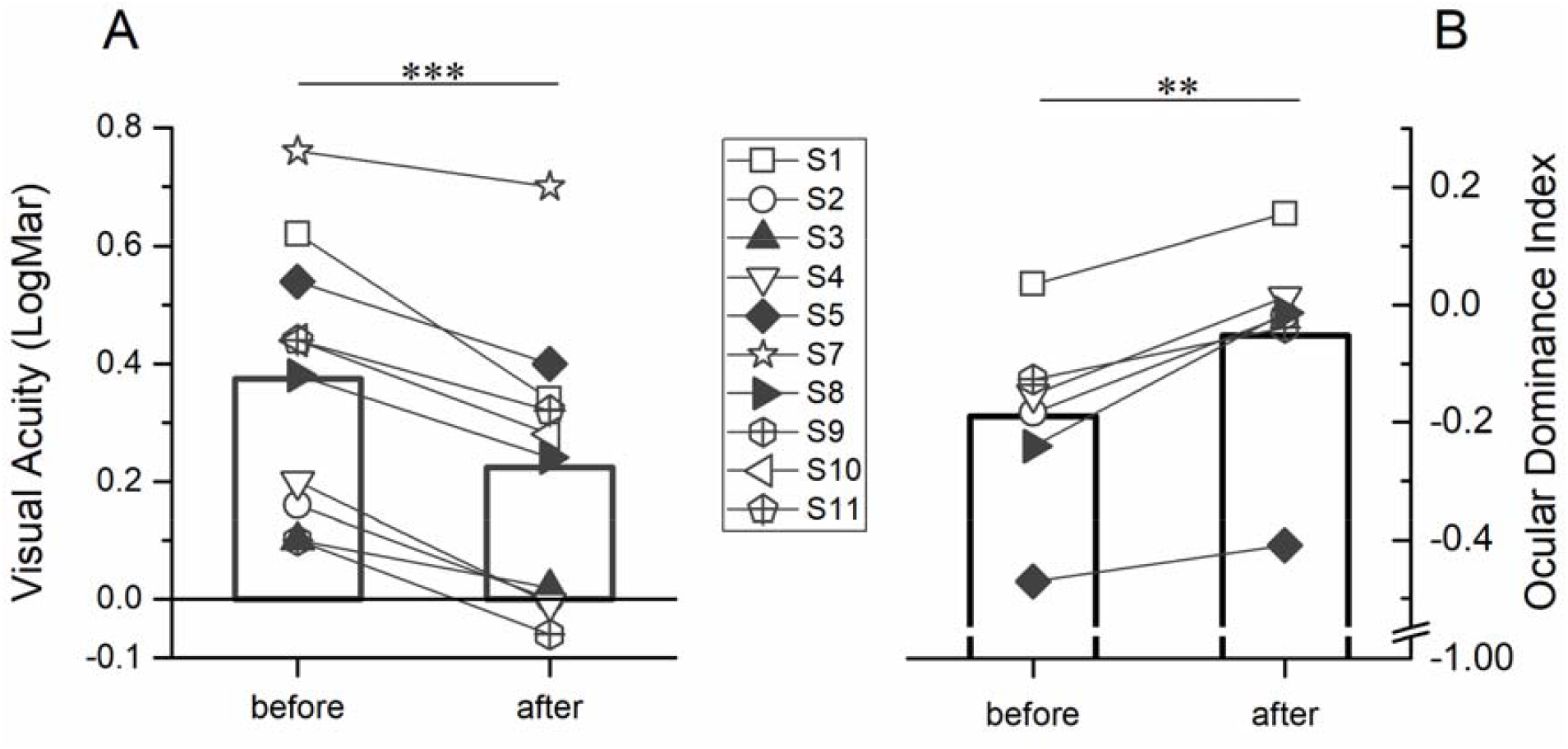
Visual acuity improvement and sensory eye dominance change after training. (A) LogMar visual acuity measured before and after the four-week training is reported for each subject (different symbols for single subjects, bars: average visual acuity). Visual acuity significantly improved after training (paired-samples t-test, *** = p<0.001). (B) Sensory eye dominance assessed before and after each 2h training session (amblyopic eye occlusion + physical exercise), averaged for each subject across training sessions (different symbols for single subjects, bars: average eye dominance). Sensory eye dominance is estimated as the contrast between the mean phase duration of the amblyopic (deprived) and non-amblyopic (non-deprived) eye in a binocular rivalry paradigm (Eq.1). The contrast of the rivalrous stimuli was adjusted before the training for each subject in order to observe perceptual alternations and eye dominance as balanced (close to the value 0) as possible (maximum contrast difference between the eyes: 100 – 20 %, amblyopic – non amblyopic eye). Even after contrast adjustment, 4/10 patients did not perceive alternations in visual dominance during binocular rivalry (complete dominance of the non-amblyopic eye). Dominance of the amblyopic eye increased after short-term monocular deprivation combined with physical exercise (paired-samples t-test, ** = p<0.001).

**Figure 3.**
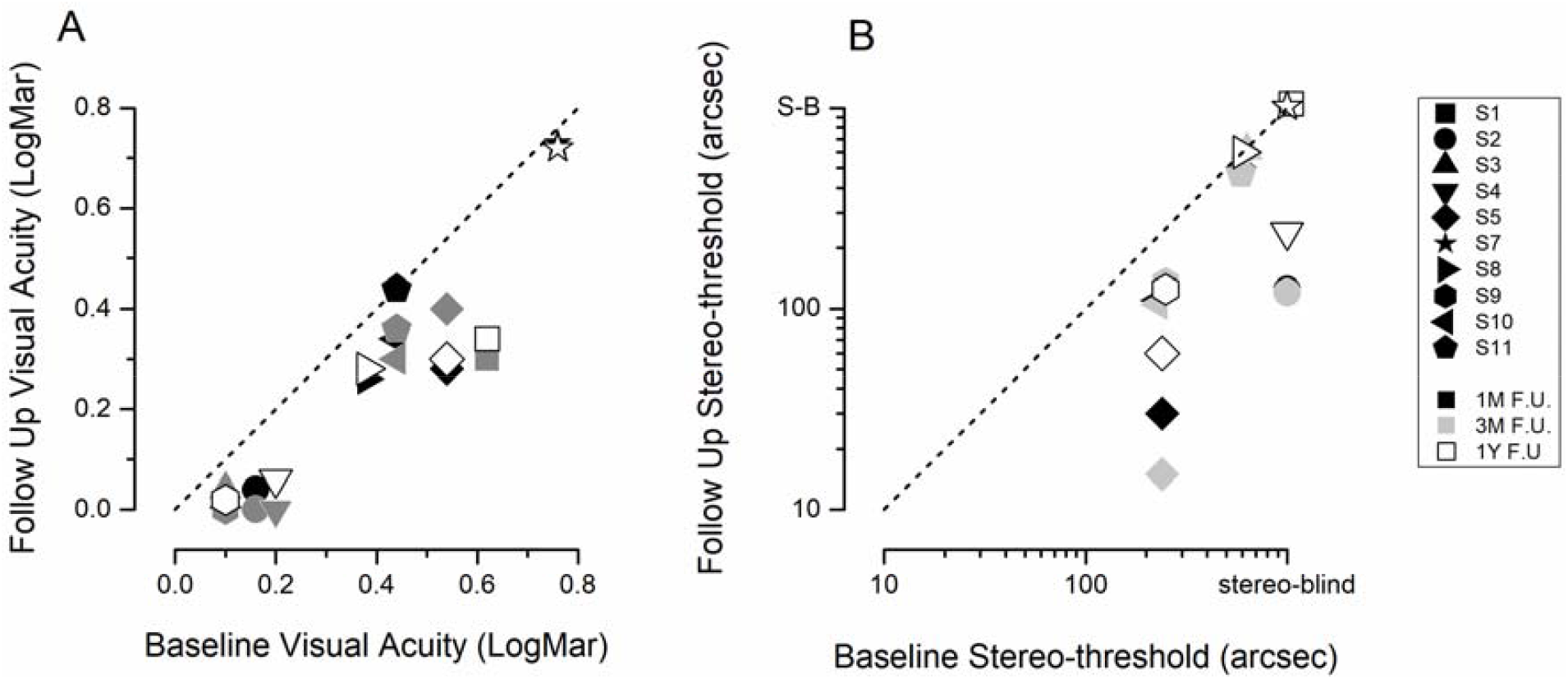
Visual acuity and stereo-threshold in follow-up measurements. (A) Scatter plot of LogMar visual acuity measured before training (x axis) and in follow-up measurements obtained one month (black symbols), three months (grey symbols) and one year (white symbols) after the end of training (different symbols shapes represent different patients). All points lie under the equality line indicating that the visual acuity improvement was maintained in follow-up measurements. (B) Same as (A) but for stereo-thresholds: patients who improved stereo-vision during the training maintained the improvement in follow-up measurements.

**Figure 4.**
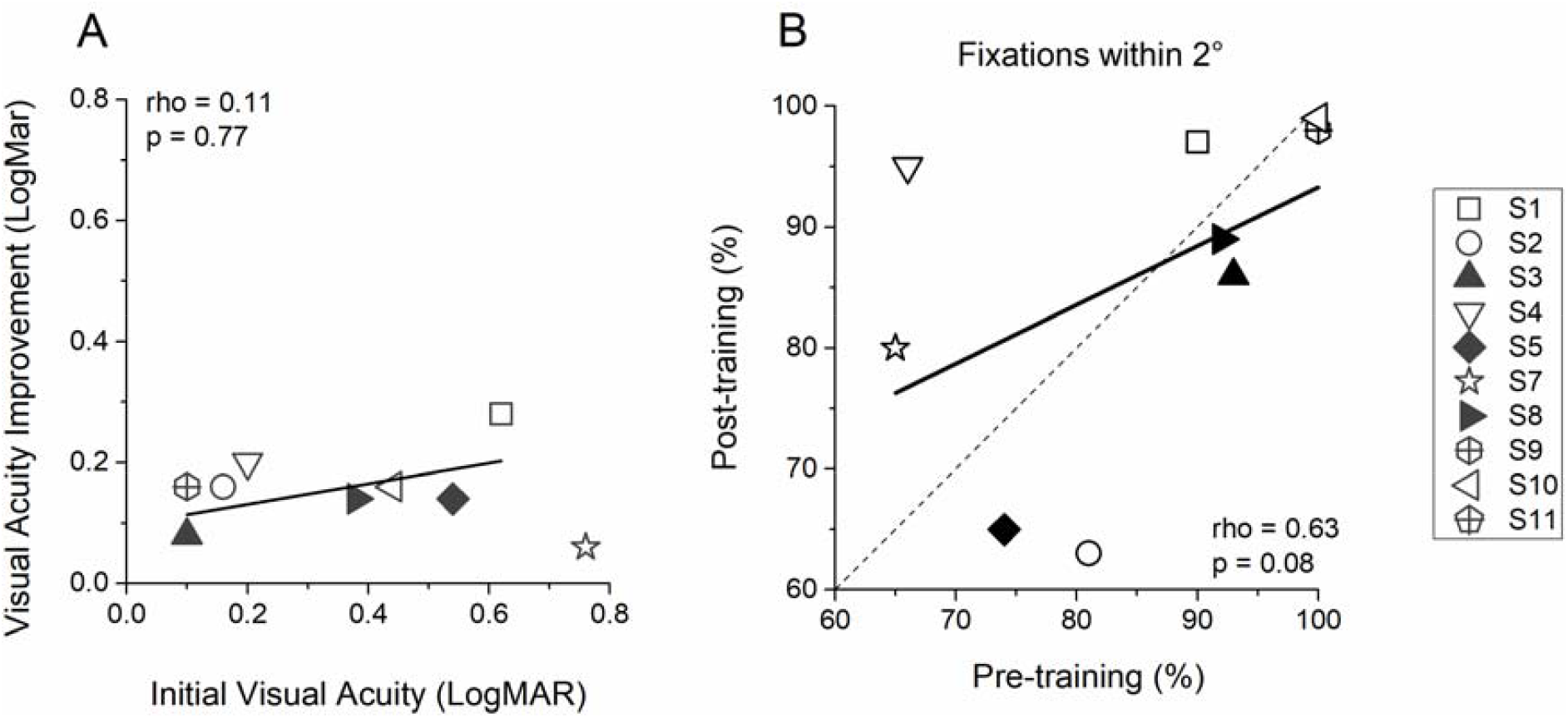
Correlations between anisometropia, visual acuity improvement and fixation quality. (A) Scatter plot of the LogMar visual acuity measured for each subject (different symbols) before training (x axis) and the difference between visual acuity measured after and before the 4-week training (y axis). No correlation is observed between initial anisometropia and visual acuity. (B) Scatter plot reporting the percentage of fixations falling within a 2-degree radius from the fixation cross presented inside a microperimeter obtained before and after training. No consistent change in fixation quality is observed across subjects.

The training was also effective enough to induce a significant improvement of stereo-thresholds (TNO test) in six out of the ten patients, independently of their occlusion therapy history (Figure 1C)., This was a significant improvement for the sample (Repeated Measures ANOVA, F(6, 54)=6.2, p=0.02, η^2^=0.7). Two of these patients were completely stereoblind before training (Figure 1C). The patients who improved stereopsis during training also maintained the improvement in the follow-up measurements up to one year (Figure 3B). Fixation quality of the amblyopic eye did not change after training (Figure 4B, fixations within 2° pre-training = 84.6±4.8%, after training=85.8±4.9%, t(8)=0.26, p=0.8), indicating that the improvement visual function was not attributable to a change in ocular motility. Interestingly, the sensory dominance of the amblyopic eye, measured with a binocular rivalry task, also significantly increased after short-term occlusion (Figure 2B, t(5)=5.26, p=0.003), suggesting a change in the binocular neuronal circuitry. The post-training changes in visual acuity, stereo-acuity and ocular dominance did not correlate with each other (all ps>0.77).

In debriefing questionnaires, all patients reported a qualitative improvement of vision in the amblyopic eye (“I have a sharper/more contrasted/brighter vision from the amblyopic eye”). Two patients also reported a significant reduction in the occurrence of headaches/migraines usually experienced after prolonged exposure to screens, three patients reported an improvement in depth perception and one reported a reduction in crowding effects (“before the training I used to confuse adjacent letters with each other”).

To disentangle the contribution of the two experimental manipulations (eye-patching and physical exercise), five of the ten patients also performed a control experiment, at least one month before being engaged in the main experiment with physical training. During the control sessions, the amblyopic eye was patched for two hours in three consecutive days without simultaneous physical exercise, but with a visual stimulation totally equivalent to that of the main experimental condition (movie). Visual acuity was found to slightly improve in the control experiment (average visual acuity improvement: 0.06±0.01 LogMar, Repeated Measures ANOVA: F(3,12)=11.18, p=0.005, η^2^=0.74, Figure7A). However, the visual acuity improvement achieved during the first three sessions of the main condition combining monocular deprivation and physical exercise was about double (Bootstrap sign-test, 10^6^ repetitions, p<0.001, Figure 5C) than that in the control experiment (average visual acuity improvement 0.1±0.01 LogMar, Figure 5C). Finally, in the control condition stereo-thresholds remained unchanged in all patients (Figure 5B), indicating that physical exercise might be crucial to enhance visual plasticity and promote the recovery of visual function in adult amblyopes.

**Figure 5.**
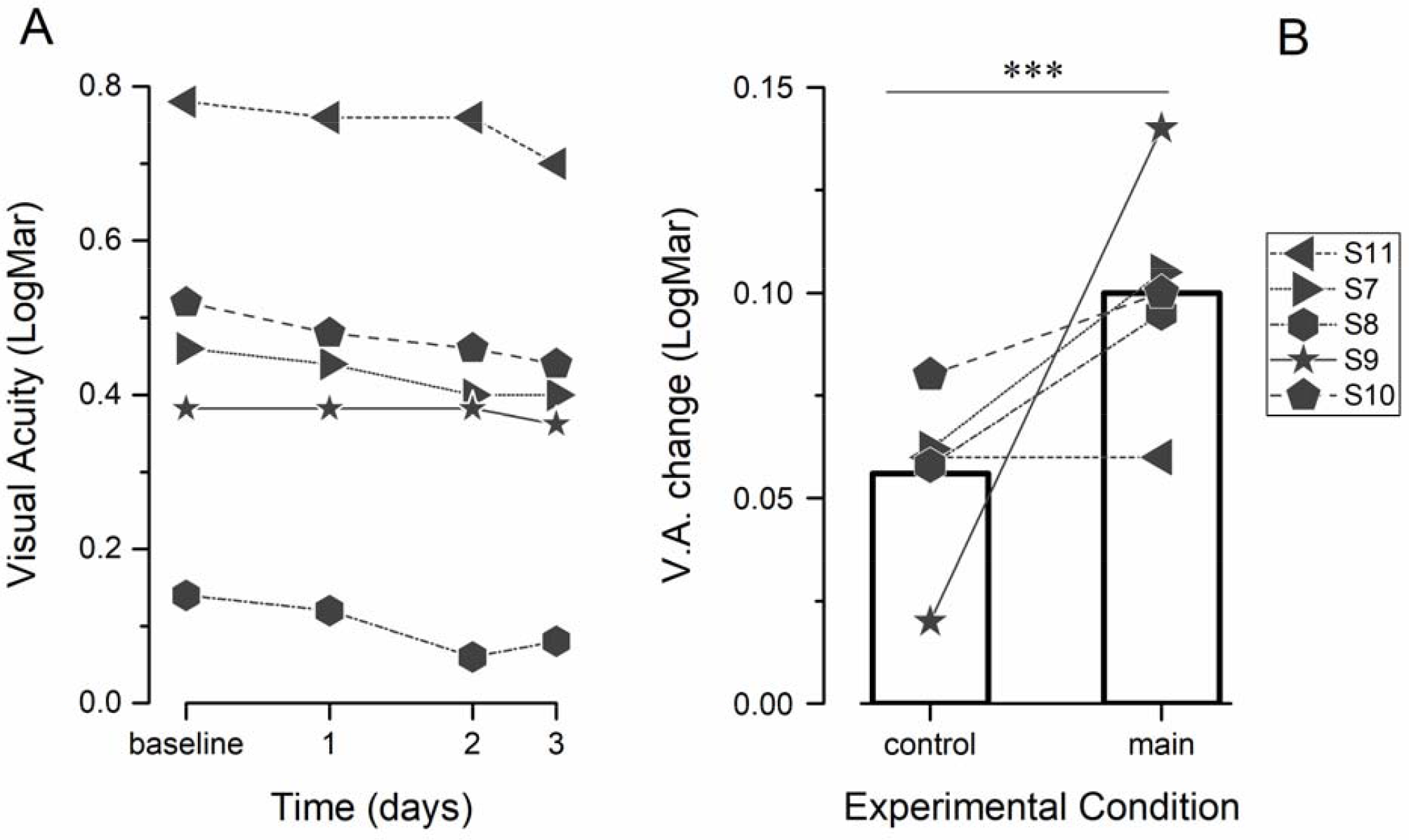
Control experiment results. Five out of the 10 patients also performed a control experiment (at least one month before taking part in the main experiment). In three consecutive days, the non-amblyopic eye was patched without the simultaneous physical exercise performance. (A) LogMar visual acuity plotted as a function of time from the beginning of the control experiment (different symbols represent individual patient performance). Only a small improvement in visual acuity was observed in the control experiment. (B) LogMar stereo-threshold plotted as a function of time from the beginning of the control experiment. Stereo-thresholds did not change in the control experiment. (C) Comparison between the visual acuity improvement observed in the control experiment and after the first three days of training in the main experiment combining monocular occlusion and physical exercise (different symbols for single subjects, bars: average visual acuity). The visual acuity improvement is significantly larger (bootstrap sign-test, *** = p<0.001) for the main experiment.

## Discussion

Together, our results show that six brief sessions of short-term deprivation of the amblyopic eye combined with physical activity promote a long-term recovery of both visual acuity and stereopsis in adult anisometropic patients. The first surprising aspect of our results is that patching the amblyopic eye is effective in improving visual acuity in anisometropic patients, a procedure that is opposite to the traditional occlusion therapy used for amblyopia^3^. Patching the amblyopic eye is commonly referred to as “inverse occlusion”^32^ and has been historically used as an alternative treatment for cases of amblyopia with eccentric fixation, aimed at preventing the reinforcement of the eccentric fixation point in the amblyopic eye^33–36^. Even though prolonged inverse occlusion (12 days) was found to be effective to some extent in this particular class of patients, its effect was weaker than the effect of traditional occlusion^36^, and this practice was abandoned. Our results show that short-term inverse occlusion (combined with physical exercise) might be more effective (up to 20 times faster) than traditional occlusion for the recovery of visual function in adult amblyopes. Moreover, the effects of short-term inverse occlusion is not due to a change in ocular motility, as fixation quality of the amblyopic eye did not change after training. Rather, we propose that the effect is due to a genuine boost of the amblyopic eye signal, consistent with the homeostatic plasticity previously observed in adult emmetropes^19–22,37,38^ and amblyopes^39,40^.

Our data show that moderate physical exercise boosts the effect of short-term monocular deprivation, inducing a larger improvement of visual acuity and stereo-sensitivity compared to inverse occlusion alone, and might be crucial to promote the long-term effect of the training. Our findings are consistent with recent evidence that voluntary physical exercise enhances visual cortical activity^41^ and promotes visual plasticity in adult rodents^27,29^ and in normally sighted humans^26,42,43^. Our results also show that physical exercise boosts visual cortical plasticity in amblyopic human subjects, after the closure of the critical period. The modulation of visual cortical activity and plasticity by physical activity has been linked to a decrease of GABAergic inhibition in the primary visual cortex of mice and rats^28,30^. We have recently found that a decrease of GABAergic inhibition in the primary visual cortex is also one of the key mechanisms mediating the effect of short-term monocular deprivation in adult humans: GABA concentration decreases in V1 after 2h30 of monocular deprivation and the change in GABA strongly correlates with the change in ocular dominance measured with binocular rivalry^37^. It is therefore plausible that the beneficial effect of exercise on visual plasticity observed here is mediated by a modulation of GABAergic inhibition, promoting ocular dominance plasticity. More generally, physical exercise also increases neurotrophic factors (BDNF, IGF-1 and VEGF) and cardiovascular fitness, two factors that might be involved in mediating neuroplasticity^44^. Increased BDNF following exercise has been related to enhanced hippocampal plasticity and neurogenesis^45,46^, as well as with improved memory and executive functions in humans^47^. Importantly, BDNF is also one of the critical mechanisms underlying visual plasticity, regulating the critical period for ocular dominance plasticity^48–50^. On the other hand, whereas cardiovascular fitness has been related to higher cognitive performances, no consistent correlation between cardiovascular fitness and improved cognitive functions after physical exercise has been found^51^ and there is no evidence at present of a role of cardiovascular fitness in mediating visual cortical plasticity.

In conclusion, we propose a new training paradigm that is totally non-invasive, does not require extensive supervision, and leaves the patients free to perform pleasant activities such as watching a movie or television. It is therefore a valid candidate for clinical applications. Amblyopia is the principal cause of (predominantly monocular) visual loss in the pediatric population (prevalence up to 5%^3^). While we believe that our data can provide a valid therapeutic strategy for adult amblyopic patients and adolescents resilient to the standard occlusion therapy, we are well aware that the methods may be not appropriate as a substitution of standard clinical therapy, especially in young children. Homeostatic plasticity of the patched eye has been reported in amblyopic children^39^ suggesting that the same method in principle could be used, but given the high plasticity of young children visual cortex it should be first carefully validated with a dose-dependent chart.

## Acknowledgements

The authors are grateful to Dr. Roberto Caputo for help with the selection of patients and continual advice, Drs. Aris Dendramis and Anna Bettinelli for help during the data collection and Prof. David Burr for helpful comments throughout the project.

## Authors Contributions

C.L., A.S. and M.C.M. designed the experiment; A.T.S., A.L., M.L and D.L performed patients’ clinical examinations; C.L. collected and analyzed the data; all authors discussed the results; C.L., M.C.M., A.T.S. and A.S. wrote the paper.

